# Multi-omics characterization of SIRT3 metabolism and its adaptation to the presence of amyloid-beta oligomers in nasal epithelial cells

**DOI:** 10.64898/2026.03.13.711520

**Authors:** Paz Cartas-Cejudo, Marina De Miguel, Silvia Romero-Murillo, Andrea A Felizardo-Otalvaro, Leire Extramiana, Joaquín Fernández-Irigoyen, Enrique Santamaria

**Affiliations:** Clinical Neuroproteomics Unit, Navarrabiomed, Hospital Universitario de Navarra (HUN), Biochemistry Area, Department of Health Science, Universidad Pública de Navarra (UPNA), Navarra Institute for Health Research (IdiSNA), Pamplona, Spain

**Keywords:** Sirtuins, SIRT3, Amyloid-beta, Alzheimer, multi-omics, sex

## Abstract

Sirtuins (SIRTs) are nicotinamide adenine dinucleotide (NAD□)-dependent deacetylases that regulate cellular homeostasis in a multifactorial manner. Although alterations in SIRT signaling are evidenced in both olfactory dysfunction and Alzheimer’s disease (AD), the specific role of sirtuin 3 (SIRT3) in olfactory metabolism remains unknown. Here, we have evidenced a partial interdependency between SIRT3 and SIRT5 deacetylase members in human nasal epithelial cell cultures (hNECs). A multi-omic integrative approach applied to conditions of SIRT3 silencing or overexpression revealed that hNEC metabolism is markedly more sensitive to reduced SIRT3 levels, identifying specific transcripts and phosphorylation sites belonging to inflammatory and redox mediators that are tightly regulated by SIRT3 in hNECs. Following exposure to oligomeric Aβ peptide, phosphoproteomic alterations promoted an activation trend of stress-induced senescence and apoptotic signaling in SIRT3-silenced hNECs, whereas induced activation of mitotic phase–related pathways, Hippo signaling, and glycogen metabolism were evidenced in SIRT3-overexpressing hNECs. From a translational point of view, a dissimilar sex-dependent profile in serum SIRT protein levels (SIRT1 and SIRT6) was observed across multiple neurological disorders including AD, mixed dementia, frontotemporal lobar degeneration and amyotrophic lateral sclerosis. These data shed new light on novel SIRT-dependent mechanisms associated with neurodegeneration, underscoring that the maintenance of optimal SIRT3 protein levels may partially counteract the detrimental effects induced by Aβ oligomers in AD at olfactory level.

## INTRODUCTION

Olfactory dysfunction is a common feature in Alzheimer’s disease (AD) [1], being considered as a premature sign of neurodegeneration and consequently, a reliable marker [2]. The olfactory epithelium-bulb (OB)-tract (OT) axis is the central site for the processing of olfactory information in the brain [3–5]. At olfactory epithelium level, amyloid-beta (Aβ) deposition can predict mild cognitive impairment and is associated with dementia [6] and AD pathology correlates with brain pathology [7]. A massive analysis has revealed that the presence and severity of neuropathology in OB-OT axis reflects the presence and severity of AD in other brain regions [7]. A significant degeneration of axons has been detected in the OT from AD subjects [8]. In pre-AD subjects, the loss of fiber OT integrity corresponds to a loss of gray matter density in parallel with a reduced glucose metabolism [9,10]. Interestingly, OTs undergo early and sequential morphological alterations that correlate with the development of dementia [11]. All these data together with experimental evidence obtained in human AD and different animal models [12,13] point out that the nasal cavity may be a useful site for early AD diagnosis [14] as well as for innovative intranasal therapies [15].

Our group has previously characterized the molecular alterations that accompany the OT neuropathological deposits in AD. This proteome dyshomeostasis induced a disruption in OT protein interaction networks and widespread sex-dependent pathway perturbations, among them Sirtuin (SIRT) signaling [16]. SIRTs are 7 class III histone deacetylase enzymes (SIRT1-7) with unique enzymatic activities, subcellular locations and substrate scopes [17,18] that present anti-inflammatory, antiapoptotic and neuroprotective effects [19]. Specifically, SIRT3, the primary mitochondrial deacetylase, displayed a tangled protein expression profile across the olfactory-entorhinal-amygdaloid axis in AD, evidencing a down regulation at the level of OT and entorhinal cortex [16]. SIRT3 regulates mitochondrial homeostasis through the reversible deacetylation of target proteins involved in antioxidant response, OXPHOS, the TCA cycle and fatty acid oxidation, being postulated as a redox-sensitive interface between cellular environment and metabolism [20,21]. However, the role of SIRT3 and its crosstalk with neuropathological insults at olfactory level remains largely unknown. Our study aims to increase our knowledge about the role of SIRT3 in olfactory metabolism, elucidating its potential interactions in AD-related contexts. To achieve this goal, we have combined silencing and overexpression strategies with olfactory *in vitro* studies as well as RNA sequencing (RNA-seq), quantitative proteomics and phosphoproteomics coupled to bioinformatics analyses. In addition, serum SIRT levels have been monitored across several human neurodegenerative diseases to extend our knowledge about SIRT metabolism and neurodegeneration.

## MATERIALS AND METHODS

### Cell culture

Human nasal epithelial cells (hNECs) (ABM. T9243) were maintained in DMEM (Gibco, Grand Island, NY, USA, 41966029) supplemented with 10% FBS (Merck Millipore, Burlington, MA, USA, A3A5256801) and 1% penicillin/streptomycin (ABM, P4333-100ML). Cultures were kept in a humidified atmosphere containing 5% CO□ at 37 °C. Cells were transfected with SIRT3 siRNA (siSIRT3, MBS827374, Sino Biological) or a negative control siRNA (5 nmol, Dismed, MBS8241404) using Opti-MEM reduced-serum medium (Gibco, 15392402). siRNA complexes were prepared according to the manufacturer’s instructions and added to the cells for the indicated incubation period before downstream analyses. For SIRT3 overexpression, hNECs were seeded in 6-well plates at approximately 80% confluency. Briefly, 1 µg of either SIRT3 (HG13033-UT, Sino Biological) or control GFP (CV026, Sino Biological) plasmid DNA was mixed with Lipofectamine 3000 (Fisher, Waltham, MA, USA, L3000015) at a 1:2 ratio in DMEM and incubated for 15 min at room temperature. The preparation is adjusted to a final concentration of 1 µM when added to the cultures. Aβ_1–42_ peptide (Sigma, Burlington, MA, USA, 4014447) was resuspended in anhydrous DMSO (Thermo Fisher Scientific, D12345), sonicated for ten minutes, supplemented with DMEM, and stored at 4 °C for 24 hours. The preparation was adjusted to a final concentration of 1 µM when added to the cultures. Twenty-four hours post-transfection, Aβ_1–42_ oligomer (1 µM) was added. First, after an additional twenty-four hours incubation, total RNA was isolated using the RNeasy Mini Kit (Qiagen, Hilden, Germany, 74106) according to the manufacturer’s protocol.

### RNA sequencing (RNA-seq) and Data Analysis

Total RNA was extracted and purified using the RNeasy Mini Kit (Qiagen, Hilden, Germany) according to the manufacturer’s protocol. Sequencing libraries were generated using the Illumina Stranded Total RNA Prep with Ribo-Zero Plus kit (Illumina Inc., San Diego, CA, USA) from 100 ng of total RNA, following the depletion procedure as instructed. Libraries were sequenced on a HiSeq1500 system (Illumina) in PE100 rapid mode and pooled in equimolar proportions to a final concentration of 10 nM. Library concentrations were quantified using a Qubit 3.0 fluorometer (Invitrogen, Waltham, MA, USA), and fragment size distributions were verified by capillary electrophoresis on a Fragment Analyzer (AATI). RNA-seq data quality was first assessed using FastQC v0.11.9 (http://www.bioinformatics.babraham.ac.uk/projects/fastqc/) and MultiQC v1.9 (http://multiqc.info/). Raw reads were trimmed and filtered to retain sequences with a Phred quality score ≥25, and adapter sequences were removed using TrimGalore v0.5.0 (https://www.bioinformatics.babraham.ac.uk/projects/trim_galore/). To eliminate residual rRNA reads, trimmed sequences were processed with SortMeRNA v2.1 (https://bioinfo.lifl.fr/RNA/sortmerna/) [22] to remove 18S and 28S rRNA fragments that might remain after depletion. High-quality reads were aligned to the Homo sapiens reference genome (GRCm38.p6/GCA_000001635.8, ftp://ftp.ensembl.org) using HISAT2 v2.2.1 (https://daehwankimlab.github.io/hisat2/) [23] under default parameters. Alignment quality was evaluated with Qualimap2 (http://qualimap.bioinfo.cipf.es) [24], and resulting BAM files were sorted and indexed using Samtools v1.18 [25]. Gene-level read counts were obtained with FeatureCounts (http://subread.sourceforge.net) [25,26], and differential expression analysis between experimental conditions was conducted with edgeR [27]. To streamline and automate group comparisons, the SARTools pipeline was implemented [28]. All output data were compiled into HTML reports and CSV tables containing count density plots, pairwise scatter plots, clustering dendrograms, principal component analyses (PCoA), size factor estimations, dispersion plots, and MA/Volcano plots. The resulting CSV file, including raw and normalized counts, fold-change estimates, and dispersion metrics, was annotated with additional gene information from the Biomart database (https://www.ensembl.org/biomart/martview/346d6d487e88676fd509a1b9a642edb2). To control the false discovery rate (FDR), p-values were adjusted using the Benjamini–Hochberg (BH) correction method. Genes with adjusted p-values <0.05 and fold-change values >1.3 or <0.7 were considered significantly up- or down-regulated, respectively. (All websites were accessed on February 14, 2024.).

### Sample preparation for shotgun proteomics

Before processing, all samples were randomized, and experimental groups were balanced to minimize potential bias during handling. Cell pellets were homogenized in a lysis buffer containing 8 M urea (Sigma, Burlington, MA, USA), 50 mM dithiothreitol (DTT) (Merck, 43815), and benzonase (Merck, E1014), supplemented with protease (cOmplete Mini, Roche #11836153001) and phosphatase inhibitors (PhosSTOP, Roche #4906845001). Lysates were centrifuged at 20,000 x g for 1 h at 15 °C, and the resulting supernatant was quantified using the Bradford assay kit (Bio-Rad, Barcelona, Spain #5000006). For phosphopeptide analysis, 600 µg of total protein was subjected to enzymatic digestion. Proteins were reduced with DTT (final concentration 20 mM, 30 min at room temperature), alkylated with iodoacetamide (Merck, I1149)(final concentration 30 mM, 30 min at room temperature in the dark), diluted to 0.9 M with ammonium bicarbonate (ABC) (Merck, A6141), and digested with trypsin (Promega, Madison, WI, USA, V5111) at a 1:20 (w/w) enzyme-to-protein ratio for 18 h at 37 °C. Digestion was stopped by acidification (pH < 6, acetic acid), and peptides were desalted using Pierce™ Peptide Desalting Spin Columns (Thermo Fisher Scientific, Waltham, MA, USA, 89852). Phosphopeptide enrichment was performed with the High-Select™ TiO_2_ Phosphopeptide Enrichment Kit (Thermo Fisher Scientific, A32993) following the manufacturer’s protocol. The enriched phosphopeptide fraction was then cleaned as described above and dried using a SpeedVac concentrator. A 10-µg aliquot of desalted peptides from the total protein digest was reserved for global proteome analysis.

### Data independent acquisition (DIA)-mass spectrometry

Dried peptide samples were reconstituted in 2% acetonitrile with 0.1% formic acid (ACN-FA), spiked with internal retention time (iRT) peptide standards (Biognosys), and quantified using a NanoDrop™ spectrophotometer (Thermo Fisher Scientific) prior to LC-MS/MS analysis. Peptide separation was performed on an EASY-nLC 1000 system coupled to an Orbitrap Exploris 480 mass spectrometer (Thermo Fisher Scientific). Samples were loaded onto a C18 Aurora column (75 µm x 25 cm, 1.6 µm particles; IonOpticks) and eluted at 300 nL/min using a 60-min gradient at 50 °C as follows: 2-5% buffer B in 1 min, 5-20% B in 48 min, 20-32% B in 12 min, and 32-95% B in 1 min (buffer A: 0.1% FA; buffer B: 100% ACN with 0.1% FA). Ionization was achieved with a spray voltage of 1.6 kV and a capillary temperature of 275 °C. Data were acquired in data-independent acquisition (DIA) mode with full MS scans over 400-900 m/z (resolution: 60,000; maximum injection time: 22 ms; normalized AGC target: 300%) and 24 MS/MS maximum injection time: 22 ms; normalized AGC target: 100%). Peptide fragmentation was performed using higher-energy collisional dissociation (HCD) with a normalized collision energy of 30%.

### Bioinformatics and Statistical Analysis

Mass spectrometry raw data were processed using Spectronaut (Biognosys) through direct DIA (dDIA) analysis. MS/MS spectra were searched against the Homo sapiens UniProt reference proteome (UP000005640) using standard parameters. The enzyme specificity was set to trypsin. For total proteome analysis, carbamidomethylation (C) was defined as a fixed modification, while oxidation (M), acetylation (protein N-terminus), deamidation (N), and pyro-glutamate formation (Q) were set as variable modifications. For phosphoproteome analysis, carbamidomethylation (C) was also defined as a fixed modification, with oxidation (M), acetylation (protein N-terminus), and phosphorylation (S, T, Y) included as variable modifications. Protein and peptide identifications were filtered at a 1% Q-value (FDR) threshold. Quantitative data from total proteome analyses were exported to Perseus (version 1.6.15.0) [29] for statistical evaluation and visualization. Pairwise comparisons were performed using unpaired Student’s t-tests, with statistical significance defined as p < 0.05 and a 1% peptide-level FDR. Proteins exhibiting an absolute fold-change <0.77 or >1.3 (linear scale) were considered significantly downregulated or upregulated, respectively. Phosphoproteomic data were processes using the Peptide Collapse plugin (v1.4.4) in Perseus [30], converting Spectronaut peptide reports to site-level datasets. Default settings were applied, grouping post-translational modifications (PTMs) by sample (FileName), collapsing the data matrix at the site level, and filtering PTM localization probabilities to >0.75. Statistical analyses were performed following the same procedure as for total proteome data. To investigate the association of differentially expressed genes and proteins with dysregulated regulatory or metabolic networks, several bioinformatic tools were employed, including Metascape [31], QIAGEN’s Ingenuity Pathway Analysis (IPA) (v1.2, QIAGEN, Redwood City, CA, USA). Metascape was utilized to extract biological insights related to -ome functionality. IPA calculates the statistical significance of associations between biological or molecular events and the imported molecules using Fisher’s exact test (p ≤ 0.05). Furthermore, IPA comparison analysis hierarchically ranked signaling pathways based on the calculated p-values.

### RNA isolation and Reverse Transcription Quantitative Polymerase Chain Reaction (RT-qPCR)

For RT-qPCR experiments, total RNA from cells was extracted with Mini RNeasy Kit for RNA Purification (Qiagen, Hilden, Germany). In all cases, RNA was treated with DNAse (Invitrogen, 18068015) and reverse transcription was performed using PrimeScript RT Reagent Kit (Takara, RR037A). RT-qPCR was performed using SYBR Green (Thermo Fisher, 4309155) and a QuantStudio 12K Flex qPCR system (Applied Biosystems) with triplicate biological replicates for each sample, and fold change was calculated normalized to GAPDH expression. A list of primer sequences is provided as Supplementary table 1.

### Western blotting

Equal protein amounts (5 µg) were resolved in 4-15% stain free SDS-PAGE gels (BioRad) and transferred onto nitrocellulose membranes using the Trans-Blot Turbo transfer system (up to 25 V, 7 min; Bio-Rad). Membranes were incubated with primary antibodies (1:1000 dilution) in 5% nonfat milk or BSA, following the manufacturer’s recommendations. Anti-SIRT1 (#9475), anti-SIRT2 (#12650) and anti-SIRT3 (#5490) were from Cell Signaling. Washing, membranes were incubated with horseradish peroxidase-conjugated secondary antibodies (1:5000), and immunoreactive bands were visualized using enhanced chemiluminescence (PerkinElmer) and imaged with a ChemiDoc MP Imaging System (Bio-Rad). Equal protein loading was verified using stain-free imaging technology. Protein normalization was performed by quantifying total protein directly from the stain-free gels used for Western blotting, as previously described [32]. Densitometric analysis was carried out using Image Lab Software (version 5.2; Bio-Rad), and optical density values were expressed in arbitrary units and normalized to total protein levels within each lane.

### Human serum samples

Serum samples and data from patients included in the study (Supplementary table 1) were provided by the Biobank at CIMA-Universidad de Navarra and were processed following standard operating procedures approved by the Clinical Ethics Committee of Navarra Health Service (study code: PI_2024/3).

### Enzyme-Linked Immunosorbent Assay (ELISA)

Serum SIRT1 (MBS2601311), SIRT2 (MBS2701791), SIRT3 (MBS9714317), SIRT4 (MBS2020879), SIRT5 (MBS2705671), SIRT6 (MBS162109) and SIRT7 (MBS455811) concentrations were measured using enzyme-linked immunosorbent assay kits according to the manufacturer’s instructions (MyBioSource). Data were analyzed using GraphPad Prism Software v8. Mann-Whitney U test was used for between-group comparisons. We considered p-value less than 0.05 to be statistically significant.

## RESULTS

### Metabolic effects of SIRT3 modulation in nasal epithelial cells

We have established specific hNEC systems silencing and overexpressing SIRT3 to decipher metabolic adaptations associated to SIRT3 dynamic protein changes (Figure 1A and 1D). Both genetic modifications did not significantly modify the macroscopic phenotype of hNECs (Figure 1B and 1E). Subsequent experiments were performed to explore potential SIRT3-dependent modulations in the mRNA expression of the SIRT family. In *SIRT3* silencing conditions, we observed a significant increment in *SIRT5* mRNA levels (Figure 1C). However, no compensation changes were detected in SIRT3 overexpression conditions (Figure 1F), indicating a moderate crosstalk and a partial interdependency between SIRT deacetylase members in hNEC cultures. A multi-omic approach was used to specifically characterize molecular events associated to SIRT3 silencing or overexpression conditions as well as to identify direct functional mediators of SIRT3 in hNECs at transcriptomic, proteomic and phosphoproteomic levels. According to our statistical criteria, SIRT3 silencing induced 1716 differentially expressed genes (DEGs) (724 down / 992 up), 125 differentially expressed proteins (DEPs) (60 down / 65 up) and 388 differential phosphosites (DPPs) (148 down /240 up) respect to scrambled control condition. However, SIRT3 overexpression induced 72 DEGs (49 down / 23 up), 13 DEPs (3 down / 10 up) and 497 DPPs (275 down / 222 up) (Supplementary table 2-4). An integrative analysis detected a subset of transcripts and phosphosites that directly depends on SIRT3 levels. As shown in Figure 2A, the transcript levels corresponding to *VCAN*, *IL6*, *KIF3C*, *MARCKS*, *ARL4C*, *IL1B* and *SOX4* were increased in *SIRT3* silencing conditions whereas their levels dropped in overexpression conditions. In the same context, *IL7R*, *TXNRD1*, *NQO1* and *SLC7A11* diminished their transcriptional level in SIRT3 silencing conditions, being overexpressed in parallel with SIRT3 overproduction (Figure 2B). . At phosphoproteomic level, six phosphosites were directly dependent on SIRT3 protein levels (Figure 2B). KPNA (S6), SRRM2 (S857) and LEO1 (Y606) were hyperphosphorylated in overexpression conditions whereas SIRT3 silencing induced a hypophosphorylation in these residues. Moreover, LARP6 (S409), PPP1R13L (S110) and TANK (S225) were significantly increases in SIRT3 silencing conditions and decreased in SIRT3 overexpression ones. Ten phosphosites corresponding to a subset of proteins (AHNAK, LCOR, NPM1, GLYR1, DOCK7, MTDH, TOMM22, NUCK, LZTS2, MAP7D1) were equally modulated independent of SIRT3 levels (Figure 2B). All these data suggest that: i) hNEC transcriptome is more sensitive to a decrease in SIRT3 levels and ii) a subset of differential features (transcripts and phosphosites) corresponding to inflammatory and redox mediators are tightly regulated by SIRT3 in hNECs. Whereas biological functions associated to cell growth and migration were significantly affected at the three omics layers in SIRT3-silenced cells (Figure 2C), most of the GO terms significantly altered by the SIRT3 overexpression were governed by phosphorylation events related to mRNA metabolism and chromatin remodeling (Figure 2D). A more detailed functional analysis is also shown in supplementary figures 1 and 2, respectively.

**Figure 1:**
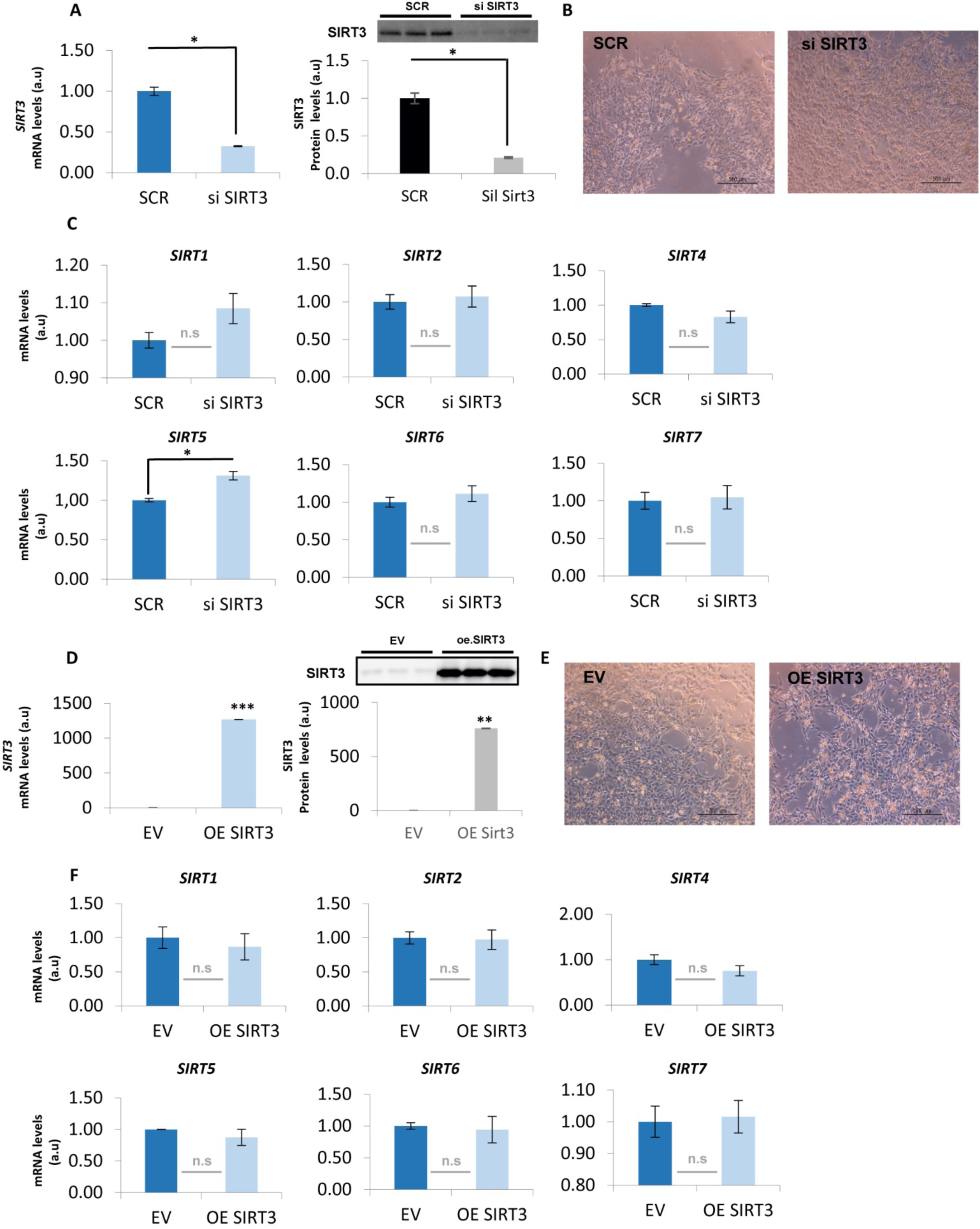
SIRT3 silencing/overexpression in hNECs. RT-qPCR and WB of SIRT3 in scramble (SCR) and si SIRT3 conditions (A). Representative images of SCR and si SIRT3 in hNE cell cultures (B). SIRT3 silencing effects on SIRT family mRNA abundance in hNECs (C). RT-qPCR and WB of empty vector (EV) and SIRT3 overexpression (OE) conditions (D). Representative images of EV and SIRT3 OE in hNE cell cultures (E). SIRT3 OE effects on SIRT family mRNA abundance in hNECs (F). Data are presented as mean±SEM . *P<0.05 vs. control group (a.u: arbitrary units).

**Figure 2:**
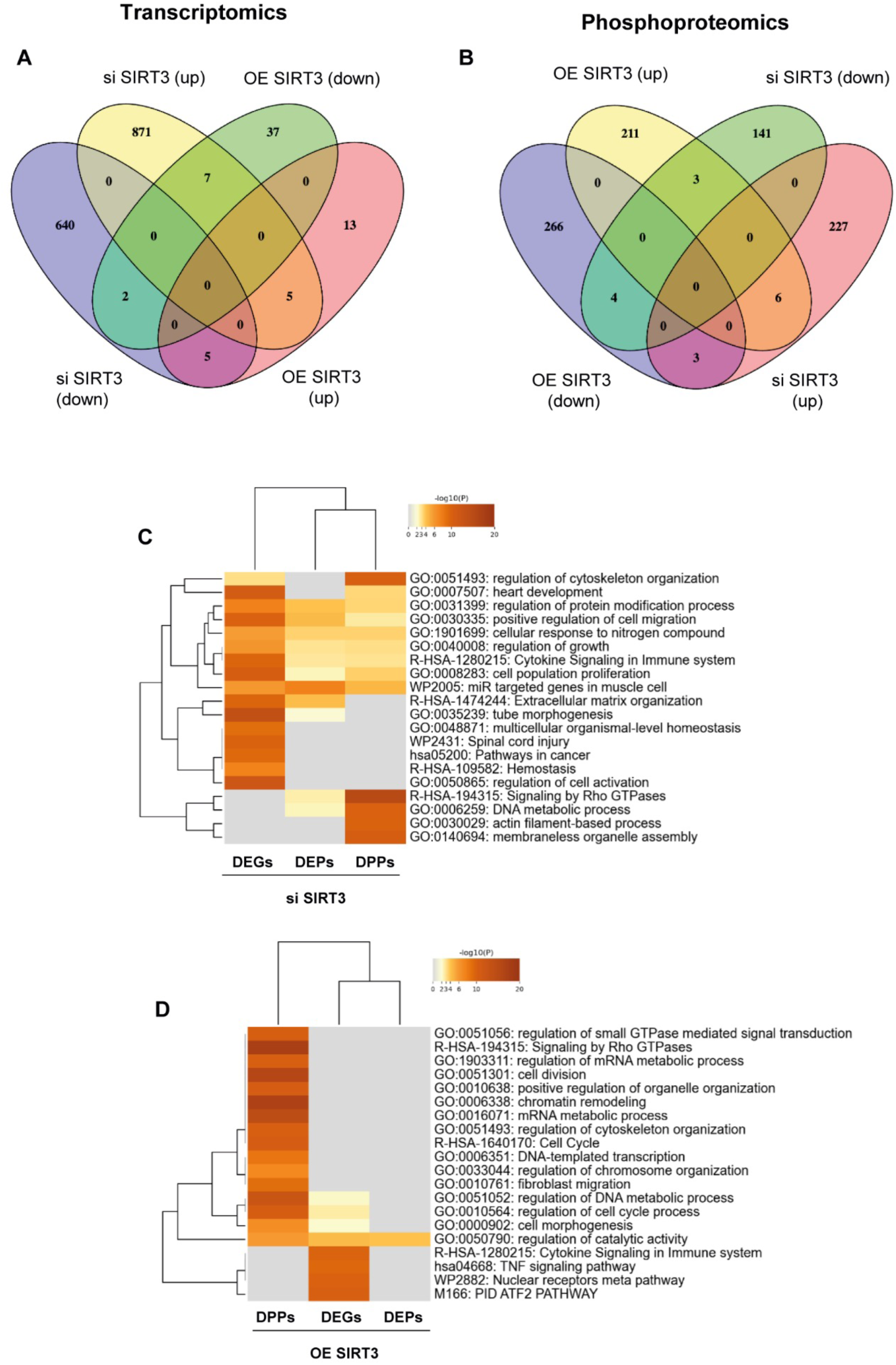
Functional analysis associated to the multi-omic approach in SIRT3 regulatory conditions. Overlap between down- and up-regulated DEGs (A) and DPPs (B) from SIRT3 silencing and overexpression conditions. Functional analysis of DEGs, DEPs and DPPs from SIRT3 silencing (si) (C) and overexpression (OE) (D).

### Metabolic effects of SIRT3 modulation in A***β*** -stimulated nasal epithelial cells

Time-dependent experiments were performed to explore the effect of Aβ (monomer/oligomer states) on SIRT protein levels in hNECs. As shown in Figure 3, SIRT1 and SIRT2 protein levels were unchanged (data not shown) whereas a significantly transient decrease in SIRT3 protein levels were observed between 8-48 hours post-treatment (Figure 3A), indicating that SIRT3 is a direct target of the substrates involved in olfactory neurodegeneration. We wanted to know the metabolic response of hNECs to the presence of Aβ oligomer in SIRT3 silencing or overexpression states. Shotgun (phospho)proteomics at 24 hours revealed: i) 34 DEPs (26 down/8 up) and 609 DPPs (354 down/255 up) in SIRT3-silenced cells treated with Aβ oligomer respect to SIRT3 silencing condition and ii) 22 DEPs (9 down/13 up) and 754 DPPs (351 down/403 up) in SIRT3-overexpressed cells treated with Aβ oligomer respect to SIRT3 overproduction condition (Supplementary tables 3 and 4 and Supporting Figure 3). These data indicate that the crosstalk between Aβ and SIRT3 is mostly mediated by phosphorylation dynamics instead of variations in protein abundance in our experimental conditions. Interestingly, we observed a SIRT3-dependent reversion of phosphorylation changes at seven p-sites (Figure 3B) corresponding to TRIM24, TBC1D9B, ZC3HC1, AKIRIN2, ZMYM2, PPP1R13L and CTNND1, most of them involved in protein turnover regulators, nuclear/chromatin dynamics, and cell□cycle/trafficking processes (Figure 3C). On the other hand, Aβ induced changes in 75 p-sites (52 down / 23 up) independent of SIRT3 levels in hNECs (Figure 3B). These phosphoproteins mainly mapped in signaling by Rho GTPases, membrane trafficking and cytoskeleton organization, pathways commonly disrupted in +/-SIRT3 Aβ-treated cell cultures (Figure 3D). Subsequent analyses were performed to monitor the SIRT3-dependent activation profile of pathways modulated by Aβ using a System-biology approach. An increase in the p53 activity by acetylation in SIRT3-silenced conditions that was reversed in SIRT3-overexpressed cells (Figure 4A), confirming the functional p53-SIRT3 axis [33] and validating our workflow. Using the IPA knowledgebase in combination with differential phosphoproteomics datasets, we observed an activating trend in stress-induced senescence, apoptosis signaling and apoptotic execution phase in SIRT3-silenced cells treated with Aβ. However, these events were partially inhibited in SIRT3-overexpressed cells upon Aβ treatment, together with a positive activation score for pathways associated to mitotic prometaphase/metaphase/anaphase, Hippo signaling, glycogen metabolism as well as clathrin-mediated endocytosis, cell junction organization and Golgi to ER traffic (Figure 4A). All these data suggest that optimal SIRT3 protein levels partially counteract the detrimental effects induced by Aβ oligomers in hNECs. Exploring the potential interlocking of SIRT3-dependent differential phosphoproteins with the amyloid-beta precursor protein (APP) metabolism, we observed that part of the DPPs detected in our study belong to the APP functional protein network (Figure 4B and 4C). The differential phosphorylation dataset associated to SIRT3-silenced cells treated with Aβ pointed out a potential inhibition of APP (Figure 4B) whereas an activating trend was evidenced for this protein in the functional network associated to SIRT3-overexpressing cells treated with Aβ (Figure 4C).

**Figure 3:**
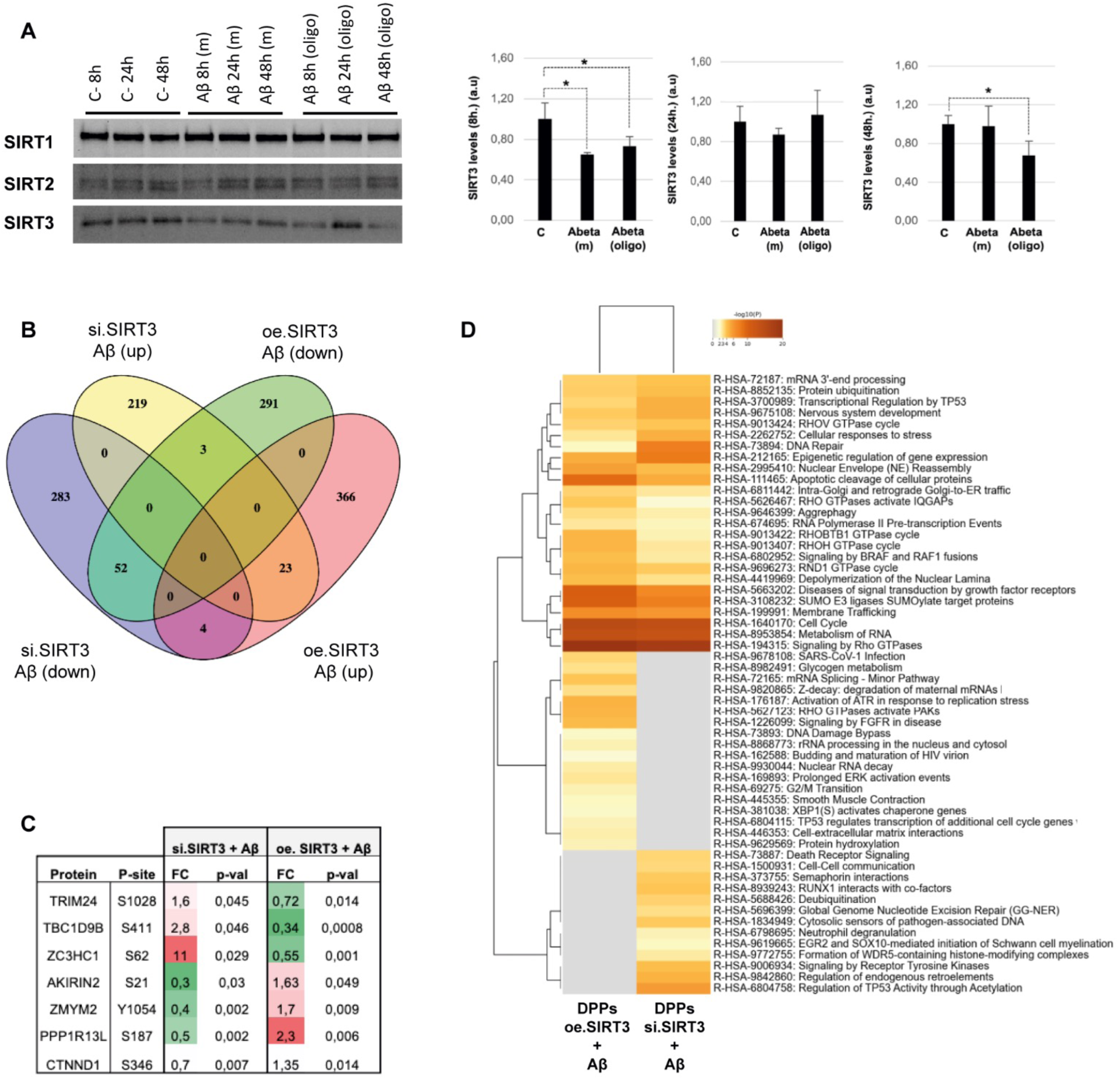
Time-dependent SIRT 1-3 protein expression variation upon Aβ (monomer/oligomer states) protein stimulation in hNECs. Steady state protein levels of SIRT 1-3 were measured by Western blotting (WB) at 8-24-48 hours post-treatment (A). * p<0.05 respect to untreated-hNEC control cells. Overlap between down- and up-regulated DPPs from SIRT3 silencing and overexpression upon Aβ oligomer protein stimulation in hNECs (B). Selected differentially phosphorylated proteins affected by SIRT3 modulation in Aβ-treated cells (C). Functional analysis of DPPs from SIRT3 silencing (si) and overexpression (OE) (D).

**Figure 4:**
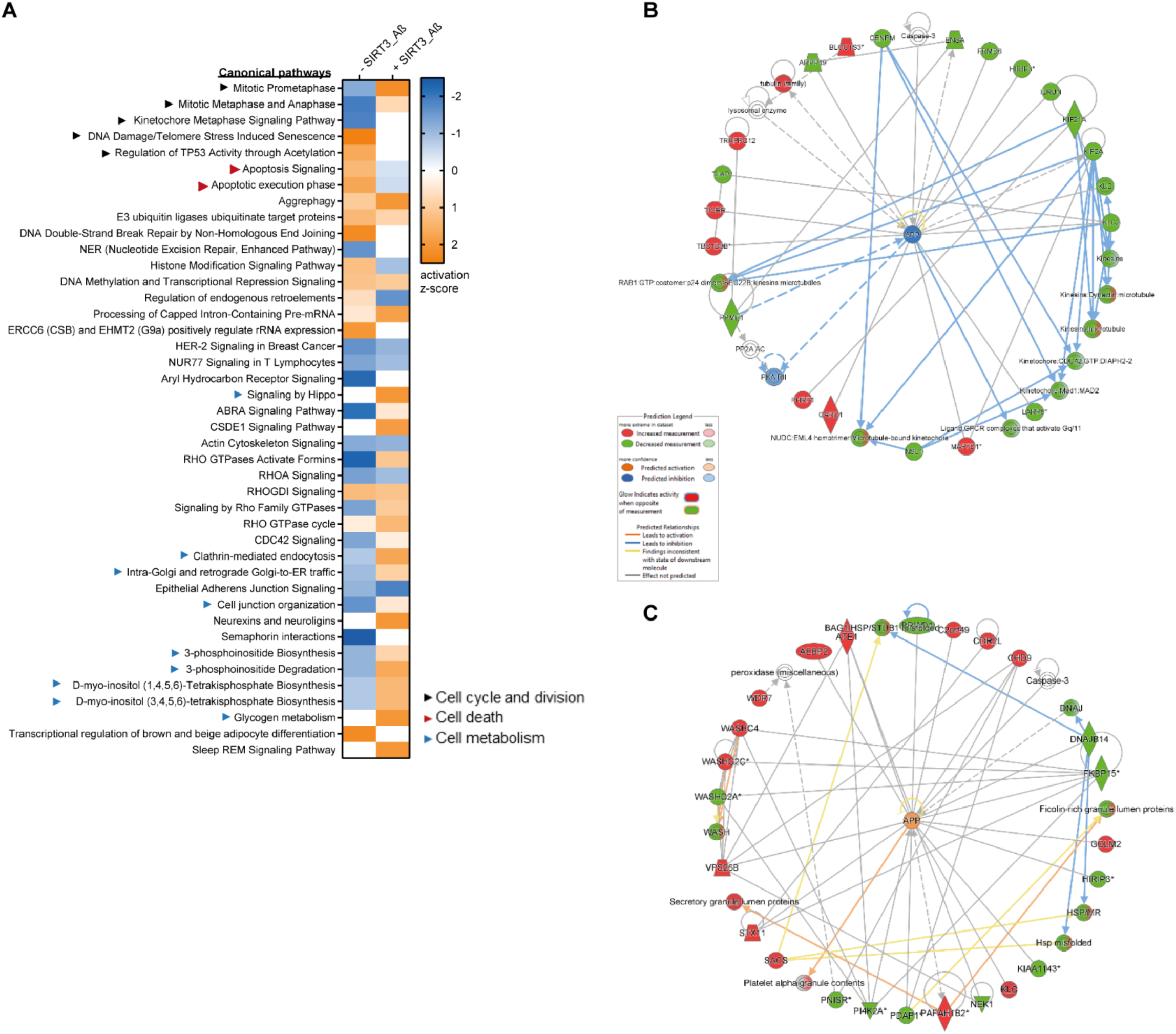
Differential signaling dynamics and phosphoprotein networks in SIRT3 silencing and overexpression conditions upon Aβ oligomer stimulation. Predictive activation profile of pathways, biofunctions and upstream regulators at the level of SIRT3 silencing and/or overexpression upon Aβ oligomer considering APP as main hub. Positive z-scores indicate potential activated pathways, whereas negative z-scores refer to predicted inhibited pathways. Based on SIRT3 silencing and overexpression datasets (A). Activation prediction of significantly altered pathways and neuronal functions (B). Systems Biology analysis were performed through the Ingenuity Pathway Analysis software [52].

### Serum SIRT protein levels across neurodegenerative diseases

Based on multiple findings on the involvement of SIRT proteins in neurodegenerative diseases, we focus our attention on the monitoring of the serum SIRT protein panel across different neurodegenerative backgrounds (See supplementary table 1). For that, all SIRT members (SIRT1-SIRT7) were evaluated in serum samples derived from controls (n=80; 40F/40M), AD (n=40; 20F/20M), mixed dementia (n=13; 8F/5M), frontotemporal lobar degeneration (FTLD) (n=13; 5F/8M) and amyotrophic lateral sclerosis (ALS) (n=12; 7F/5M). Using an ELISA approach, we only obtained quantifiable and reliable data for SIRT1 and SIRT6 at serum level. SIRT1 serum levels were significantly decreased in AD (males and females) respect the control group (Figure 5A) and SIRT6 serum levels were exclusively reduced in AD males, observing a significant difference between both sexes in the control group (Figure 5B). On the other hand, SIRT1 levels were significantly increased in mixed dementia, FTLD women and ALS backgrounds (Figure 5C). However, SIRT6 levels were exclusively reduced in FTLD men, maintaining normal levels in mixed dementia and ALS (Figure 5D).

**Figure 5:**
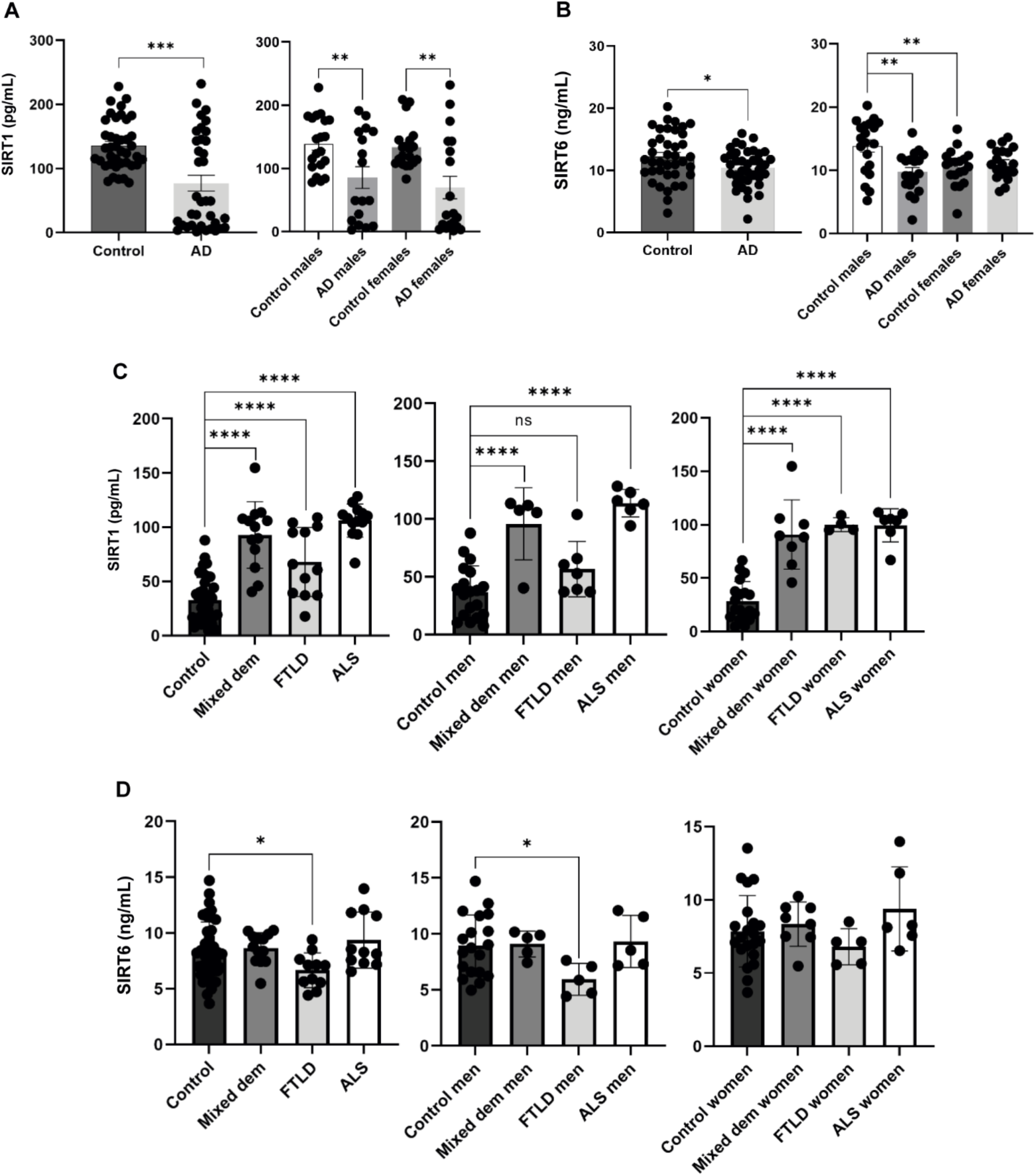
Serum SIRT family levels across neurological disorders. SIRT1 (A) and SIRT6 (B) levels measured in the serum derived from control and AD individuals, also discerning by sex. Serum SIRT6 levels from control and AD individuals, distinguishing by sex (B). Sex-specific analysis of serum SIRT1 (C) and SIRT6 (D) levels across neurological disorders. Data are presented as mean±SEM. *P<0.05 vs. control group; **P<0.01 vs. control group; ***P<0.001 vs. control group; ****P<0.0001 vs. control group (a.u: arbitrary units).

## DISCUSSION

Sirtuins are a conserved family of NAD□-dependent deacetylases that regulate key aspects of cellular physiology. However, their role in olfactory metabolism is understudied. It has been demonstrated that the olfactory function recovery after excitotoxic lesion of OBs is directly associated with increases in bulbar SIRT expression [34]. On the other hand, olfactory impairment due to perinatal protein malnutrition induces an OB SIRT downregulation [35] . Based on these findings, maintaining adequate SIRT levels seems essential for the proper preservation of olfactory function. We have previously observed a tangled SIRT protein expression profile across the olfactory-entorhinal-amygdaloid axis in AD, evidencing a SIRT3 down regulation at the level of OT and entorhinal cortex [16]. In this study, we have mapped molecular changes associated to SIRT3 modulation in hNECs in the absence or presence of Aβ to increase our knowledge about the role of this protein and its potential crosstalk with one of the neuropathological substrates associated to AD at olfactory level.

At basal level, we observed a significant increment in *SIRT5* mRNA levels when SIRT3 was silenced in hNECs. This may be due as a potential compensation mechanism, considering that both SIRTs share subcellular location and targets [36]. According to previous reports [37], our multi-omic approach clearly evidenced that cellular metabolism is more sensitive to a SIRT3 reduction in hNECs. Mechanistically, we have revealed novel mediators that interfere with the SIRT3 pathway, highlighting inflammation-associated sensors such as IL6, IL1B and IL7R. Interestingly, the olfactory epithelium functions not only as a specialized sensory interface but also as an active component of the nasal immune defense system. Chronic inflammation of the nasal mucosa is strongly associated with olfactory dysfunction, and elevated concentrations of IL-6 have been shown to compromise the barrier integrity of nasal epithelial cells, thereby promoting neuroinflammatory processes that culminate in olfactory impairment [38,39]. All these data indicate that the fine-tuning of the SIRT3-IL6 axis may contribute to maintain a proper olfactory function. Interestingly, the overexpression of SIRT3 was able to partially reverse the phosphorylation state of protein mediators canonically involved in nuclear functions like nuclear protein import (KPNA), mRNA splicing (SRRM2), RNA binding activity (LARP6) and p53/NFkB inhibition (PPP1R13L, TANK). However, further research is needed to elucidate the specific role of this protein subset in human smell.

We have observed a down-regulation of SIRT3 protein levels in Aβ -treated hNEC cultures. According to our data, the exposure of cultured cortical neurons to Aβ also downregulates SIRT3 levels [40], indicating that the protective role of SIRT3 against Aβ toxicity is not mediated by a neuron-specific mechanism. In addition, SIRT3 expression is consistently reduced in human AD brain tissue and in transgenic mouse models [16,40–44], pointing out that the loss of SIRT3 function may increase cell vulnerability to AD-related stressors.

In cerebral contexts, it has been mechanistically demonstrated that the SIRT3 protective effect may be achieved by the suppression of the BACE1-mediated amyloidogenic processing to slowdown Aβ production [37], the upregulation of Aβ-degrading enzymes [45], the attenuation of Aβ-induced neuronal hypometabolism [46], the modulation of the Wnt/β-catenin signaling pathway [47] and the prevention of p53-induced mitochondrial dysfunction [33]. In hNECs, we have observed that SIRT3 has the capacity to reshape additional signaling landscapes directly involved in the apoptotic/proliferative equilibrium, modulating the phosphorylation dynamics of the stress-induced senescence and apoptotic execution phase as well as Hippo pathway and mitotic phases in the presence of Aβ, between other routes. All these data reinforce the notion that the restoring of SIRT3 expression by natural compounds or NAD+ precursors may be beneficial to delay or prevent neurodegenerative disorders [48].

Interestingly, we have detected signaling mediators whose phosphorylation profile is tightly regulated by SIRT3 levels in the presence of Aβ in hNECs such as TRIM24, ZMYM2 (both involved in transcriptional control), TBC1D9B (GTPase activator), ZC3HC1 (component of SCF-type E3 ligase complex), AKIRIN2 (involved in nuclear protein degradation) and CTNND1 (involved in cell-cell adhesion mechanisms). These data reveal novel crosstalk between SIRT3 biology and protein phosphorylation, complementing its best-studied relationships with other posttranslational modifications like acetylation [48,49]. However, further studies are needed to understand the widespread mechanistic actions induced by the crosstalk between acetylation and phosphorylation and its role during the olfactory neurodegenerative process.

In this study, SIRT3 levels were not properly detected in human serum by an ELISA based method. However, it is important to note that all SIRT protein family has been previously detected by a more sensitive surface plasmon resonance (SPR) technique. Using the biosensor BIAcore-3000 system, Pradhan *et al.* showed that serum SIRT1, SIRT3 and SIRT6 levels were also significantly reduced in AD respect to mild cognitive impairment and control groups, proposing that SIRT members could be considered potential protein markers of AD [50]. On the other hand and in line with our observations obtained in the mixed dementia group, *Gulmammadli et al.* detected higher serum SIRT1 levels in dementia patients [51], suggesting that the type and severity of dementia may influence circulatory SIRT levels. Based on this information and the sex-dependent serum SIRT fluctuations observed in our pilot study across controls and different neurological syndromes such as FTLD and ALS, broader studies are required not only to clarify the potential value of the SIRT family as biomarkers in brain pathology, but also to monitor their levels across different stages of each neurological disorder and their potential relationships with clinical variables. Such studies are needed to determine whether alterations in SIRT proteins are more closely related to the causal origins of the pathology or represent downstream consequences of the neurodegenerative process.

## Author contributions

Conceptualization: Enrique Santamaría; data curation: Paz Cartas-Cejudo, Marina De Miguel, Joaquín Fernández-Irigoyen, and Enrique Santamaría; formal analysis: Paz Cartas□Cejudo, Marina De Miguel, Leire Extramiana, Silvia Romero-Murillo, Andrea A Felizardo-Otalvaro, Joaquín Fernández□Irigoyen, Enrique Santamaria; *in vitro* studies: Paz Cartas-Cejudo, Marina De Miguel, Leire Extramiana, Silvia Romero-Murillo, Andrea A Felizardo-Otalvaro; funding acquisition: Enrique Santamaría, Joaquín Fernández-Irigoyen; investigation: Paz Cartas□Cejudo, Marina De Miguel, Leire Extramiana, Silvia Romero-Murillo, Joaquín Fernández□Irigoyen, Enrique Santamaria; methodology: Paz Cartas□Cejudo, Marina De Miguel, Leire Extramiana, Silvia Romero-Murillo, Andrea A Felizardo-Otalvaro, Joaquín Fernández□Irigoyen, Enrique Santamaria; writing—original draft: Enrique Santamaría; and all authors gave final approval of the manuscript and are accountable for all aspects of the work.

## Acknowledgements

We are indebted to the Biobank at CIMA-Universidad de Navarra for providing us with the serum samples as well as the associated clinical and pathological data. Authors thank all JPOST Team for helping with the mass spectrometric data deposit in ProteomeXChange/PRIDE and GEO deposit. The Clinical Neuroproteomics Unit of Navarrabiomed is a member of the Spanish Olfactory Network (ROE) (supported by grant RED2022-134081-T funded by Spanish Ministry of Science and Innovation).

## Data availability statement

Mass-spectrometry data and search results files were deposited in the Proteome Xchange Consortium via the JPOST partner repository (https://repository.jpostdb.org) [53] with the identifier PXD074550 for ProteomeXchange and JPST004406 for jPOST (for reviewers: https://repository.jpostdb.org/preview/40335519869948d4851f70 Access key: 9563). The RNA-sequencing data generated in this study have been deposited in the Gene Expression Omnibus (GEO) under accession number GSE324335.

## Ethics statement

According to the Declaration of Helsinki, all assessments and experimental procedures were previously approved by the Clinical Ethics Committee of Navarra Health Service (study code: PI_2024/3).

---

**Supplementary Figure 1:**
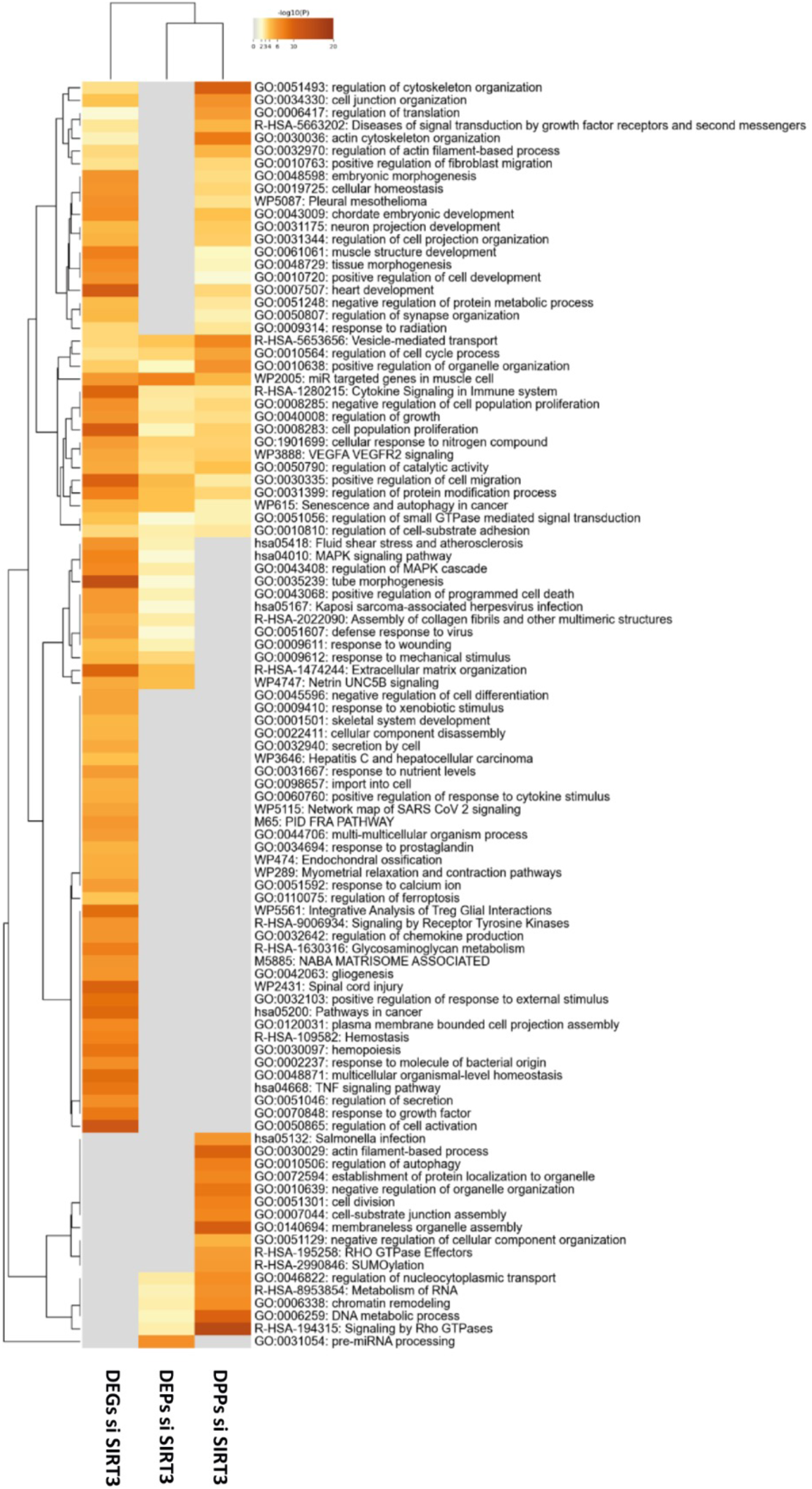
Functional mapping of deregulated genes (DEGs), proteins (DEPs) and phosphoproteins (DPPs) upon SIRT3 silencing conditions in hNECs.

**Supplementary Figure 2:**
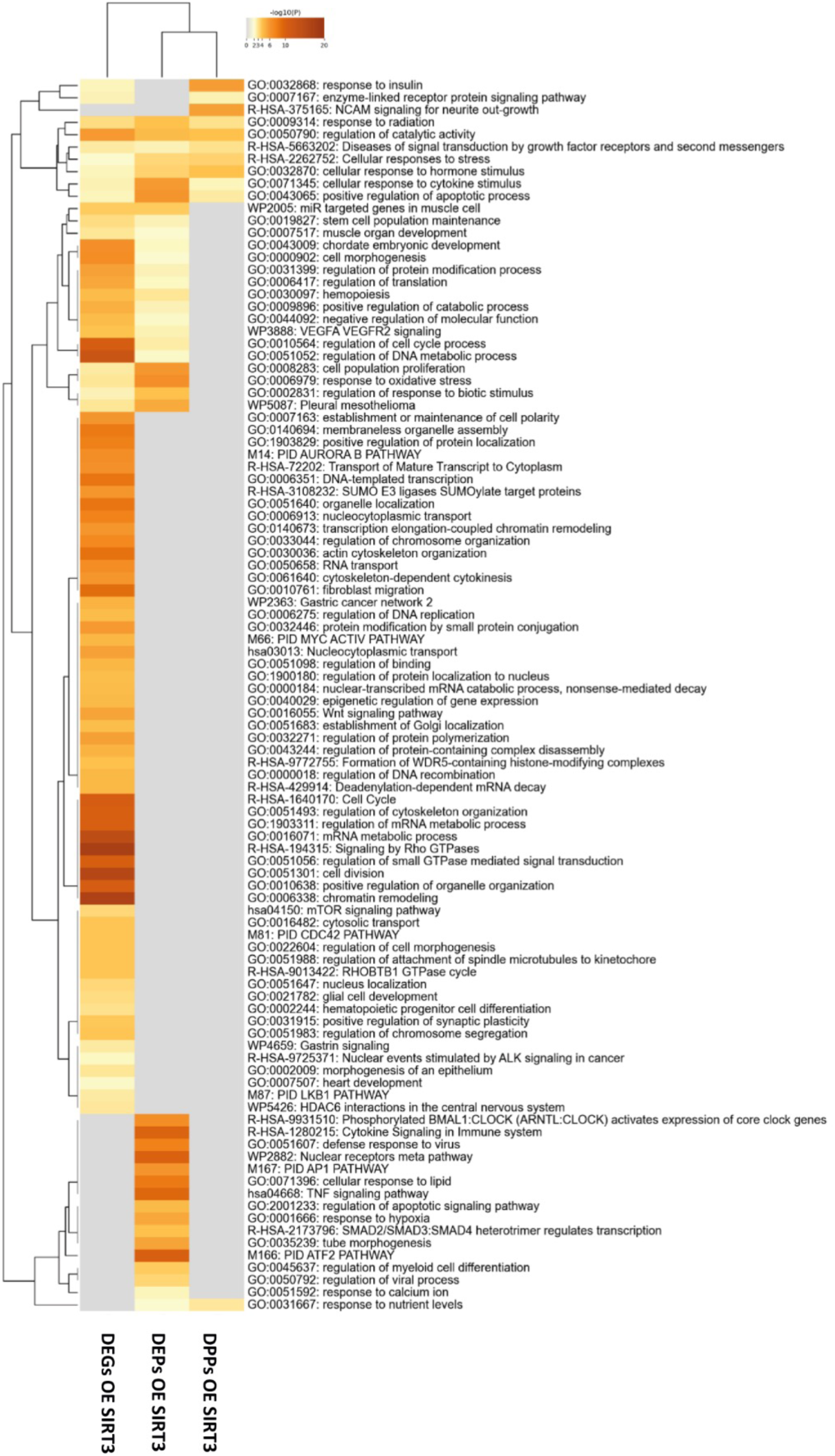
Functional mapping of deregulated genes (DEGs), proteins (DEPs) and phosphoproteins (DPPs) upon SIRT3 overexpressing conditions in hNECs.

**Supplementary Figure 3:**
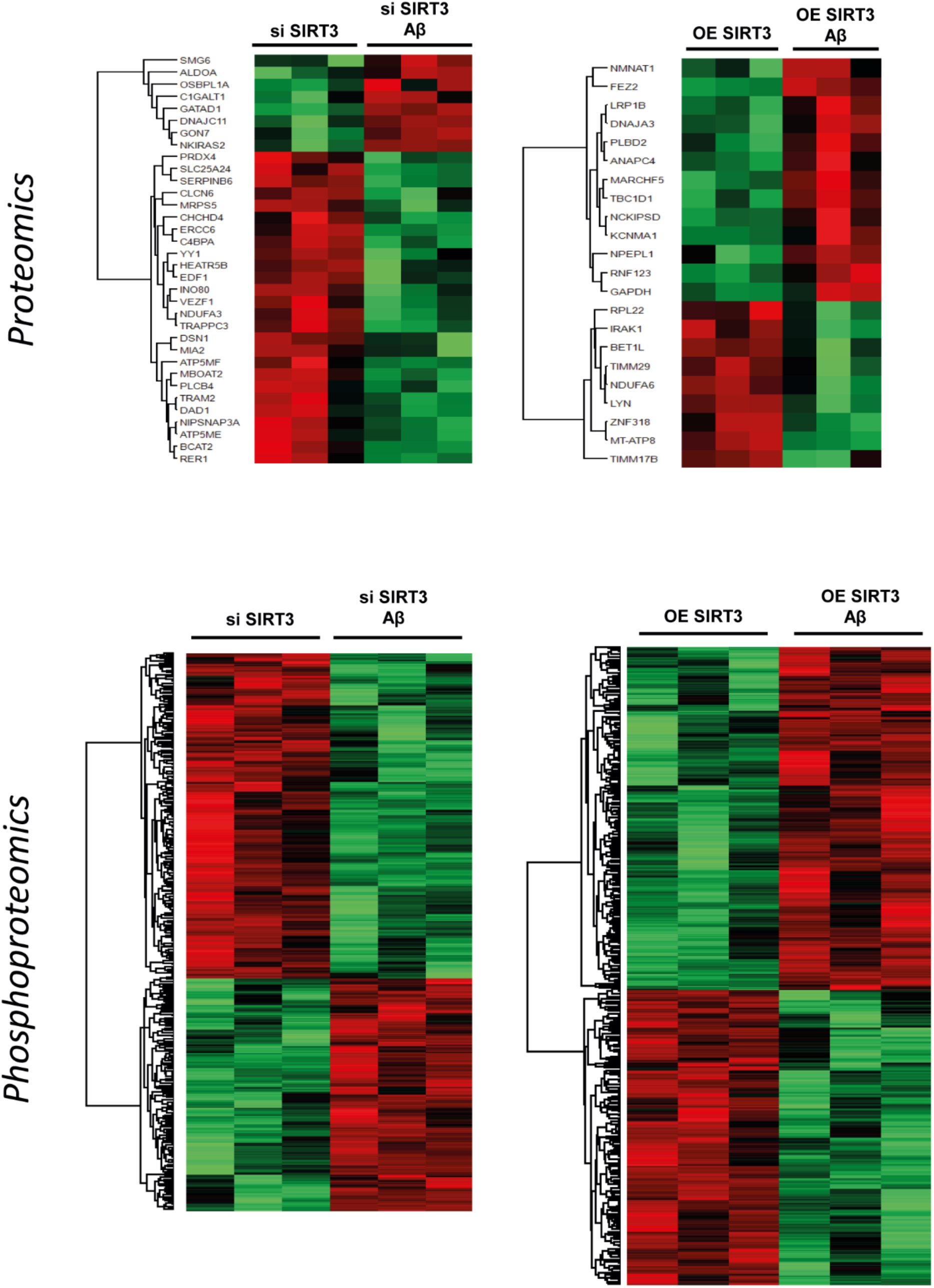
Heatmaps representing the differential proteins and phosphoproteins between SIRT3 silencing and overexpression, and upon Aβ oligomer.

**Supplementary Table 1: Primer Sequences, ELISA assays, and serum samples used in this study.**

**Supplementary Table 2: Differential genes detected upon SIRT3 silencing and overexpression in hNECs.**

**Supplementary Table 3: Significant Differential expressed proteins detected upon SIRT3 silencing and overexpression in hNECs.**

**Supplementary Table 4: Significant hypophosphorylation and hyperphosphorylation events detected upon SIRT3 silencing and overexpression in hNECs.**

